# move2utils: a utility toolkit for the move2 ecosystem

**DOI:** 10.64898/2026.07.07.736908

**Authors:** Bart Kranstauber, Kamran Safi, Anne K. Scharf

## Abstract

1. Studying animal movement at the population scale requires a stable, modern software substrate. Within R, the legacy `move` package supplied that substrate for over a decade, but its `sp`/`rgeos` backbone has been retired. The successor package `move2` deliberately confined its scope to the data class and core movebank API functions.
2. The analytical machinery of `move`, namely dynamic Brownian-bridge utilisation distributions, the directional bivariate-Gaussian variant, corridor segmentation, and along-track thinning, was left to port to the modern `sf`/`terra` stack.
3. We present `move2utils`, an R package that completes and complements that transition. `move2utils` provides `move2`-native ports of the `move` analytical functions, preserves the original C kernels where they exist, and replaces the deprecated spatial scaffolding around them. It additionally ports some of the legacy R code to faster C kernels. `move2utils` also exposes novel outlier-detection methodology described in detail in a companion paper.
4. The package is open-source (GPL ≥ 3), is developed on the MPCDF GitLab and mirrored on GitHub for public installation, and ships with vignettes and a CI-tested check suite. We illustrate it with a worked example on real tracking data and synthetic datasets.

## 1 Introduction

The R package `move2` (Kranstauber *et al*., 2024) reimplemented the foundational `move` package (Kranstauber *et al*., 2026) on the modern `sf` /`terra` stack and replaced the retired `sp` /`rgeos` backbone. By design, `move2`’s scope was confined to the data class and core functionality of data storage, base manipulations and retrieval from `movebank.org`. The analytical aspects that were part of `move`, such as utilisation distributions, corridor segmentation, along-track thinning and UD-similarity quantification were not ported. `move2utils` now closes that gap. The package supplies `move2`-native ports of the most-used `move` analytical functions, preserving the original C kernels, adding faster C kernels for some of the legacy R code, and replacing the deprecated spatial scaffolding around them. It is also the intended home for new analytical functions operating on `move2` objects. The first is an outlier-detection methodology, summarised here and described in detail in a companion paper (Safi, 2026).

## 2 Package overview

`move2utils` is implemented in R (≥4.5.0) and depends on `move2`, `sf`, `terra`, `units`, and `rlang`; analysis-specific helpers (`dplyr`, `circular`, `geosphere`, `lwgeom`) are listed under *Suggests*. The package is released under GPL (≥3) and developed openly at https://gitlab.mpcdf.mpg.de/anenvi/r-packages/move2utils, where continuous integration runs the full check suite and builds the documentation site. All exported functions that take `move2` objects are prefixed `mt_` following the `move2` convention, accept both single- and multi-track inputs, and preserve coordinate reference systems, track identifiers, timestamps, and metadata; the multi-track dispatch contract is summarised below.

Table 1 groups the 30 exported functions by theme.

**Table 1:**
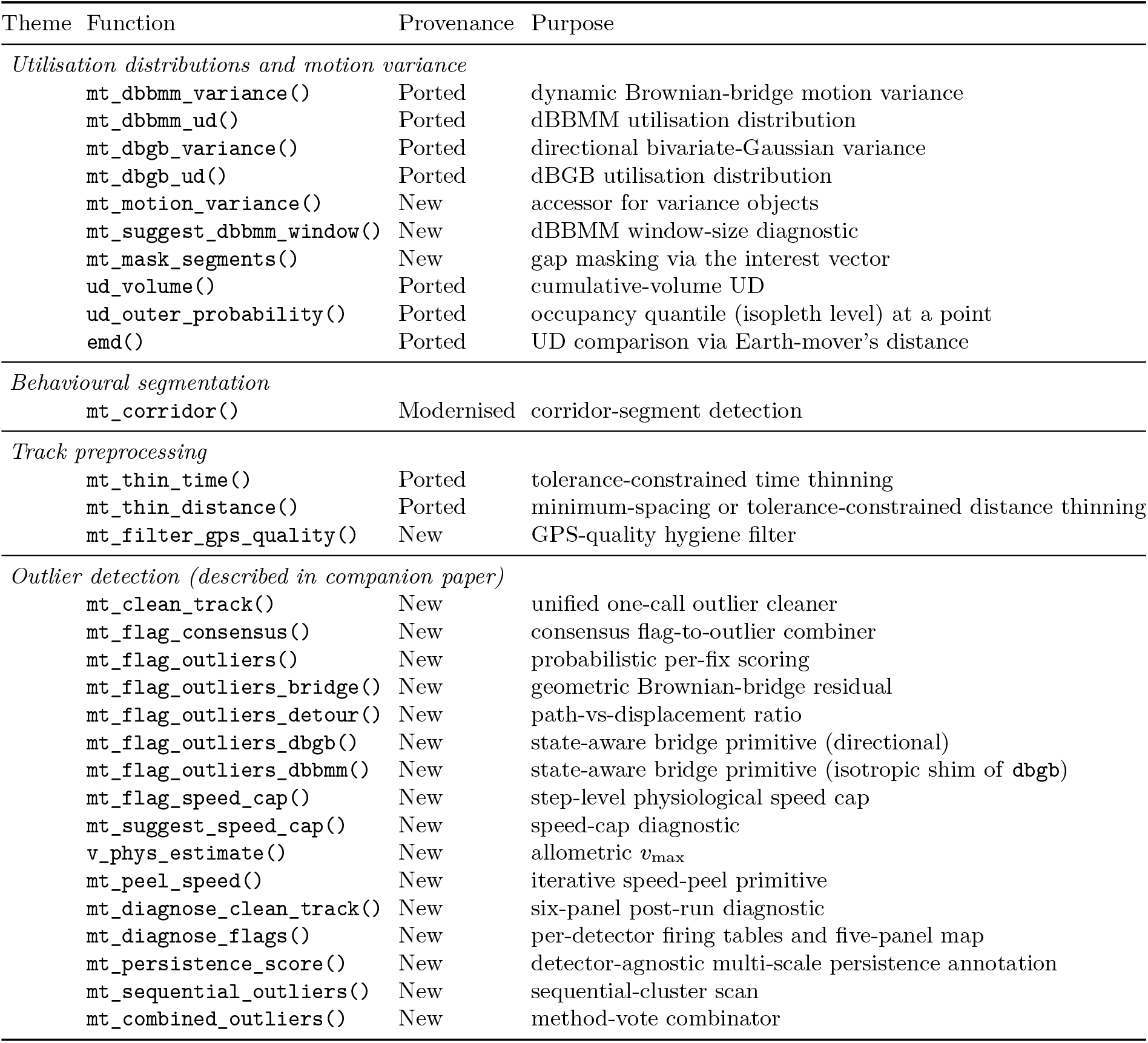
Exported functions of move2utils (v0.4.5), grouped by theme. *Ported* = adapted from a function in the `move` package, with original C kernels retained where present; *Modernised* = ported with substantial re-implementation on the `sf` /`terra` stack; *New* = first published in `move2utils`.

### Utilisation distributions and motion variance

The package’s largest theme covers dynamic Brownian-bridge motion-variance estimation and the utilisation distributions derived from it. `mt_dbbmm_variance()` and `mt_dbbmm_ud()` port the isotropic-diffusion model of Kranstauber et al. (2012) from `move`; `mt_dbgb_variance()` and `mt_dbgb_ud()` port the directional bivariate-Gaussian variant, which decomposes diffusion into the orthogonal components parallel and perpendicular to the direction of motion for trajectories with non-isotropic dispersion (Kranstauber *et al*., 2014). `mt_motion_variance()` extracts the fitted variances as a numeric vector or `data.frame`. `mt_mask_segments()` restores the gap-handling pattern of the legacy `move` workflow: on a fitted variance object it sets the `interest` flag of long temporal gaps to `FALSE`, so that `mt_dbbmm_ud()` /`mt_dbgb_ud()` drop those bridges when computing the utilisation distribution, leaving the variance estimates untouched.

Three downstream UD utilities round out the theme: `ud_volume()` converts a UD raster to cumulative-volume space for isopleth extraction, `ud_outer_probability()` returns the cumulative-volume quantile at a location — the isopleth level on which it falls, and `emd()` ports the Earth-mover’s distance of Kranstauber et al. (2017) from `move`. The Earth-mover’s distance quantifies similarity between two utilisation distributions by the minimum transport effort required to reshape one into the other, and unlike the volume-of-intersection or Bhattacharyya’s affinity it accounts for spatial proximity. The `move2utils` port keeps the simplex transportation solver of the original as a `method = “exact”` fallback, but defaults now to Sinkhorn entropic regularisation (Cuturi, 2013) together with an automated volume-based pre-mask (cells outside the `mask_quantile` contour are dropped before the cost matrix is built). The two solvers scale on different quantities: the simplex LP cost is set by the nominal grid size, whereas masked Sinkhorn scales with the *effective support* based on the cells retained inside the volume contour, so the achievable speedup depends as much on how concentrated the input UDs are as on how large the grid is. On the original `move::emd` example (`dbbmmstack`, two UDs on a 21*×*45 grid) Sinkhorn with the default 99.9 % volume pre-mask returns in ~32 ms against ~2.1 s for the simplex LP, a ~65*×* speedup at well under 1 % relative error against the LP optimum. The gap widens sharply on larger fields, where the pre-mask buys nothing: at 30*×*30 dense the LP takes ~1.9 s per pairwise call against Sinkhorn at ~95 ms (~20*×*); at 100 *×* 100 Sinkhorn returns in ~8.5 s while the simplex LP requires ~56 min on the same workstation, a ~390*×*speedup. The interface keeps `move::emd`’s `gc` and `threshold` arguments for parity. The option `gc = TRUE` switches to a Haversine ground metric for utilisation distributions on unprojected grids, and `threshold` clips the cost matrix at a maximum transport distance. The function warns when input is on a longitude/latitude grid but `gc = FALSE`, the most common silent-error pattern of the legacy function. Realistic UDs sit nearer the masked regime, so the 21 × 45 ratio is the more representative figure.

### Behavioural segmentation

`mt_corridor()` segments a track into corridor and non-corridor sections using the algorithm of LaPoint et al. (2013). The original `move::corridor()` relied on `sp` /`rgeos`; `move2utils` reimplements the spatial-indexing layer using `sf`’s R-tree, with segmentation results that match `move::corridor()` qualitatively on the same inputs.

### Track preprocessing

Two thinning helpers and one quality-control filter complement `move2`’s native filtering. `mt_thin_time()` ports `move::thinTrackTime()` and retains the strict tolerance-constrained labelling that `move2::mt_filter_per_interval()` does not provide. `mt_thin_distance()` thins by cumulative along-track distance, defaulting to a greedy minimum-spacing rule that retains one fix per `distance` of travel and offering, in parallel with `mt_thin_time()`, a tolerance-constrained mode that ports `move::thinDistanceAlongTrack()`. `mt_filter_gps_quality()` drops empty geometries and provides hooks for column-driven quality screening.

### Outlier detection

The package’s novel content is a four-primitive outlier-detection framework. `mt_flag_outliers_ bridge()` and `mt_flag_outliers_detour()` are geometric primitives. `mt_flag_outliers()` is probabilistic, leveraging a gap-aware non-parametric normalisation derived from the empirically-informed random-trajectory generator of Technitis (2021). `mt_flag_speed_cap()` is step-level and uses an allometric maximum-speed estimator based on Hirt et al. (2017). The unified entry-point `mt_clean_track()` composes these with topological block expansion and evidence-corroborated flagging, and `mt_diagnose_clean_track()` provides a six-panel post-run health check, complemented by `mt_diagnose_flags()` which returns per-detector firing tables, a consensus-comparison view, and a five-panel map for post-run audit. A `pool_by` argument on the entry-point and each primitive supports cohort-strength threshold fitting: the length-1 form pools at a single column (e.g. `“individual_id”`), and the length-2 form `c(outer, inner)` separates the fit-source level (outer) from the flag-union scope (inner) for cases such as multi-tag deployments of one animal pooled within a population. The methodology and benchmarks are presented in detail in the companion paper (Safi, 2026); we list them here for completeness.

### Design choices and performance

Two principles guided the porting. First, deprecated spatial dependencies (`sp`, `rgeos`) were replaced wholesale with `sf` /`terra`. Second, the variance-estimator C layer was extended rather than replaced: the legacy grid-evaluation kernel for dBBMM and dBGB (`src/bgb_bbmm.c`, containing the `dbbmm2()` and `bgb()` grid evaluators) was migrated from `move`, and gained a per-segment `interest` gating argument so the cascade can skip computation on segments already flagged out; an OpenMP variant (`src/bgb_bbmm_omp.c`) was added for large tracks. The dBBMM and dBGB single-window estimators were newly implemented in C (`src/bm_variance_c.c` and `src/bgb_var_window_c.c`). Each precomputes the leave-one-out quantities once per window, runs Brent’s-method 1-D optimisation on each axis, and tests every breakpoint candidate inside a single `.Call`; the dBGB version replaces an `optim()`-driven breakpoint search that the legacy `move` package ran in pure R. A track-level OpenMP outer loop wraps the per-window sweep so the C kernels saturate the available cores without R-level glue between windows. On an OUF-simulated 5 000-fix track this brings the dBGB cost from ~200*×* down to ~1.1*×* the dBBMM cost (median 0.17 s versus 0.16 s, single workstation), making the directional variant viable for the same workflows which the isotropic estimator supports. Numerical agreement with the legacy `move` implementations is asserted as a permanent regression suite (`tests/testthat/test-move-parity-*.R`): on `move::leroy` the dBBMM variance agrees with `move::brownian.motion.variance.dyn()` at a median relative difference of 1.5 *×* 10^−8^ (machine precision), the dBBMM utilisation distribution is cell-by-cell identical to `move::brownian.bridge.dyn()` at 4.4 *×*10^−9^ across ~120 000 cells, the dBGB per-axis *σ* matches `move::dynBGBvariance()` at a median absolute difference of ~5 *×* 10^−5^ m and a 99th-percentile difference below 7 cm, and the exact-LP Earth-mover’s distance matches `move::emd()` on `dbbmmstack` to within 2.5 *×* 10^−4^ relative, i.e. the LP solver’s own convergence floor. The package is gated by GitLab CI, and the cross-package parity tests are skipped on CRAN but run as part of every local check; a maintained `HEURISTICS.md` catalogues every threshold in the codebase as derived, heuristic, or empirically tuned, with rationale and plausible range.

### Working with multi-track move2 objects

The `move` package’s three trajectory classes (`Move`, `MoveStack` and `MoveBurst`) are collapsed in `move2` into a single `sf`-backed S3 class (Kranstauber *et al*., 2024). Single-track and multi-track inputs are the same object type, and track identity is recovered at call time from `mt_track_id()`; behavioural segments, which formerly required a dedicated class, are now an ordinary event-level column on the `move2` object. The shift propagates into how parameters are passed. Anything the legacy `move` user assembled into a per-individual list and dispatched by hand, e.g. per-individual measurement errors, per-burst behavioural labels, per-deployment grouping for threshold fitting, is in `move2utils` a single object-wide specification, sliced per track internally by the function.

The `location_error` argument on `mt_dbbmm_variance()`, `mt_dbgb_variance()`, `mt_flag_outliers_bridge()` and `mt_clean_track()` accepts a non-negative scalar, a length-`nrow(x)` vector, the name of a column on the object, or the literal `“auto”` (which probes `eobs_horizontal_accuracy_estimate` then `argos_lc` on the standard CLS error ladder). The `state` argument on `mt_clean_track()` accepts a column name or per-fix vector encoding behavioural states (e.g. day/night, HMM-decoded labels, season) and replaces the segmentation that `MoveBurst` used to carry, with the user owning the labels and the package owning the per-state dispatch. The `pool_by` argument on each primitive and on `mt_clean_track()` names a column (or a nested `c(outer, inner)` pair) that controls cross-track pooling of threshold-fit data. Resolved state then travels with the result: once `mt_dbbmm_variance()` slices `location_error` per track, the variance object carries the resolved per-fix vector on its `track_data`, and `mt_dbbmm_ud()` re-uses it automatically when called on the variance object or a named list of them (see Safi, 2026).

When input is multi-track the output is a named list keyed by `track_id` (`mt_dbbmm_ ud.list`, `mt_dbgb_ud.list` and `mt_mask_segments.list` dispatch on this contract); lists feed back into the next pipeline stage directly. Plotting follows the `ggplot2` and `sf` idiom — `geom_sf()` with `aes(color = mt_track_id(x))` or `facet_wrap()` on the track id — and `mt_track_lines()` provides per-track line geometries; no class-dispatched `plot()` analogue is provided. Some workflows do require an explicit per-track calculation, for instance separate utilisation-distribution grids per burst or per-sensor cleaning on a mixed-sensor tag. The idiom there is `dplyr::group_by(mt_track_id(x))` — or `dplyr::group_by(mt_track_id(x), contextAttribute)` for sub-splits — followed by the function to be applied. If the resulting object is a `move2` object, the pipe should end with `dplyr::ungroup()` to prevent downstream errors.

~~~
library(move2)
library(move2utils)
library(sf)
library(dplyr)
library(units)
fishers <- mt_read(mt_example())
fishers <- dplyr::filter(fishers, !sf::st_is_empty(fishers))
fishers <- sf::st_transform(fishers, mt_aeqd_crs(fishers))
# Multi-track input
var_all <- mt_dbbmm_variance(fishers,
                             window_size=31,
                             margin=11,
                             location_error = 20)
ud_all <- mt_dbbmm_ud(var_all, dim_size = 100) # coarse grid;
# large grids over many tracks are slow (raise dim_size for finer UDs)
# location_error stored on var_all$track_data; mt_dbbmm_ud reuses it.
# State-conditional cleaning: state lives as a column on the object.
fishers <- fishers |>
mutate(state = ifelse(mt_speed(fishers) > set_units(0.1, “m/s”),
                             “active”, “rest”))
clean <- mt_clean_track(fishers, state = “state”,
                             mass = 4.5, mode = “running”)
# Explicit iteration when settings legitimately vary per track.
var_each <- fishers |> group_by(mt_track_id(fishers)) |>
mt_dbbmm_variance(window_size = 31, margin = 11, location_error = 20)
~~~

## 3 Worked example

We illustrate a typical home-range workflow on the fisher (*Pekania pennanti*) tracking dataset bundled with `move2` via `mt_example()` (LaPoint *et al*., 2013). The chain reads a track, thins it to roughly independent fixes, fits the isotropic and directional dynamic-bridge variance models, derives a utilisation distribution from each, and identifies corridor segments, all from packaged data on a single core in under 30 s.

~~~
library(move2)
library(move2utils)
library(sf)
library(dplyr)
fishers <- mt_read(mt_example())
fishers <- dplyr::filter(fishers, !sf::st_is_empty(fishers))
leroy <- filter_track_data(fishers, .track_id = “M1”)
# remove = TRUE actually drops fixes; the default (FALSE) only marks
# retained fixes in a ‘thin_selected’ column and removes nothing.
leroy_thin <- mt_thin_time(leroy,
                             interval = as.difftime(30, units = “mins”),
                             tolerance = as.difftime(2, units = “mins”),
                             remove = TRUE)
leroy_aeqd <- sf::st_transform(leroy_thin, mt_aeqd_crs(leroy_thin))
var_iso <- mt_dbbmm_variance(leroy_aeqd,
                             window_size=31,
                             margin=11,
                             location_error = 20)
ud_iso <- mt_dbbmm_ud(var_iso)
var_dir <- mt_dbgb_variance(leroy_aeqd,
                             window_size=31,
                             margin=11,
                             location_error = 20)
ud_dir <- mt_dbgb_ud(var_dir, ext = 0.75)
# speed/circvar thresholds default to within-track quantiles (warns)
corr <- mt_corridor(leroy)
~~~

`mt_thin_time()` (called with `remove = TRUE`) reduces Leroy from 919 to 443 fixes at a thirty-minute target interval with a two-minute tolerance, suppressing burst-sampling redundancy without imposing a strict regular grid; its default `remove = FALSE` instead marks the retained fixes in a `thin_selected` column and returns the track intact. The variance estimators and UD functions accept input in any coordinate reference system and return the utilisation distribution in that same system: because the bridge geometry is computed in metres, longitude/latitude input is auto-projected to a track-local azimuthal-equidistant projection internally and the resulting raster reprojected back. The worked example above projects up front with `move2::mt_aeqd_crs(x)` onto a track-local azimuthal-equidistant grid, but this step is optional. Isopleths and area summaries are recovered with `ud_volume()` or `terra::contour()`.

Figure 1 shows the result. Panel (a) presents the thinned fisher track with the 50% and 95% UD isopleths from the isotropic dBBMM. Panel (b) overlays the directional dBGB UD for the same individual, where the elongated 50% isopleth along the principal axis of movement reflects the directional decomposition invisible to the isotropic fit. Panel (c) shows the corridor segmentation: `mt_corridor()` flags 3.0 % of fixes as corridor, concentrated along the repeated travel routes between rest sites that are characteristic of fisher home-range use.

**Figure 1:**
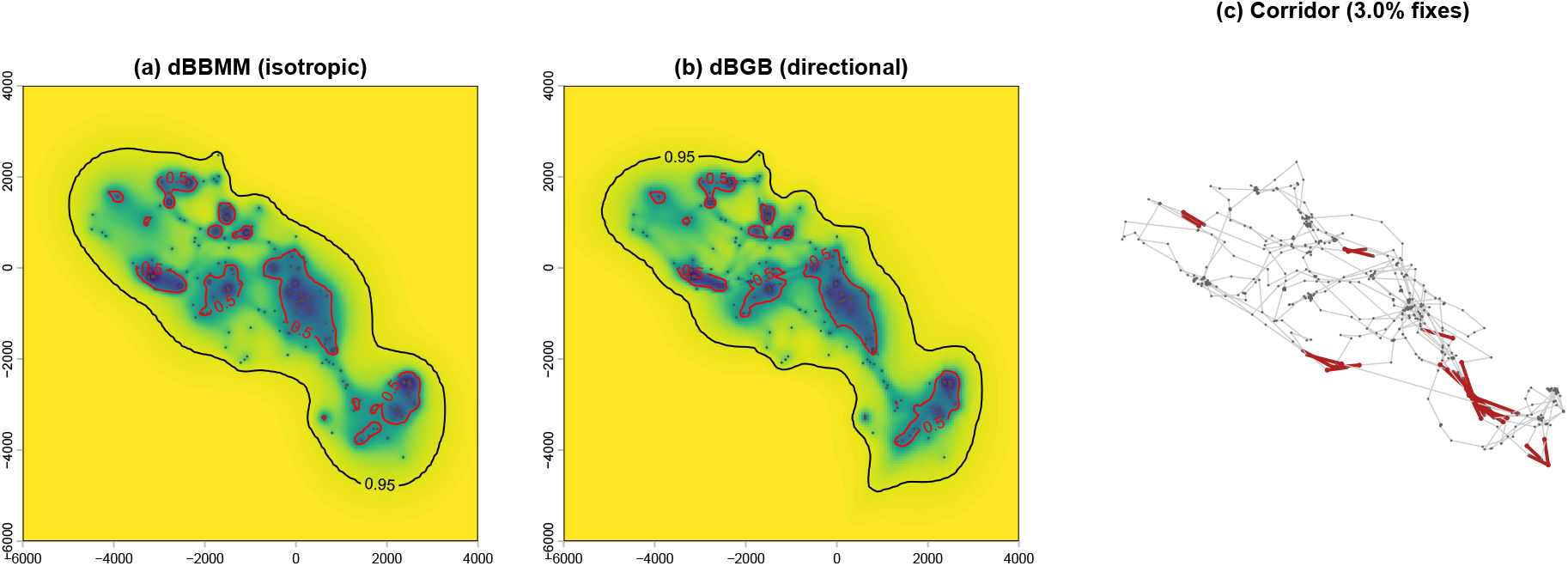
Worked example on the bundled fisher dataset (individual M1, “Leroy”; 919 raw fixes thinned to *n* = 443 at a thirty-minute target interval). (a) Thinned track with isotropic dBBMM utilisation distribution (50% and 95% isopleths). (b) Directional dBGB utilisation distribution; the 50% isopleth elongates along the principal axis of movement. (c) Corridor segmentation from `mt_corridor()`, run on the full (un-thinned) track, with the 3.0 % of fixes classified as corridor highlighted. All panels reproducible from the code listing above and the script `figures/make_figure_1.R`.

The directional UD in panel (b) carries information that the isotropic model cannot. The index of directionality *I*_*d*_ = (*σ*_*m,p*_ *− σ*_*m,o*_)*/*(*σ*_*m,p*_ + *σ*_*m,o*_), computable directly from the `mt_dbgb_variance()` object, summarises movement heterogeneity along the trajectory and can be used to elucidate changes in movement behaviour (Kranstauber *et al*., 2014). Such derived quantities are first-class outputs of `move2utils` variance objects.

Beyond home-range workflows, the same primitives underpin the outlier-detection framework. The probabilistic and bridge-residual primitives operate on the same `move2` object and compose cleanly with the variance estimators above; `mt_clean_track()` is the recommended entry-point for a single cleaning step before downstream analysis. We refer the reader to the companion paper (Safi, 2026) for the methodology and to the package’s `getting_started.Rmd` vignette for an end-to-end cleaning workflow on the same fisher data.

## 4 Conclusions

`move2utils` completes the transition of `move`’s most-used analytical machinery onto the modern `sf` /`terra` stack and consolidates that machinery into a single dependency for users adopting `move2`. The package’s scope is deliberately narrow: `move2`-native ports of established `move` functions (`ud_volume()`, `ud_outer_probability()`, `emd()` and the dBBMM/dBGB families), a small set of new utilities the porting work made natural (`mt_suggest_dbbmm_window()`, `mt_mask_segments()`, `mt_filter_gps_quality()`), and a self-contained outlier-detection framework whose methodology is the subject of a companion paper. We expect future utilities specific to movement trajectories to be absorbed into `move2utils`, providing a central place for tools operating on `move2` objects.

Within the broader R ecosystem of movement-data tools (`ctmm` : Calabrese *et al*., 2016; `amt` : Signer *et al*., 2019; `aniMotum` : Jonsen *et al*., 2023), `move2utils` occupies the niche immediately adjacent to `move2` : ports and utilities that operate directly on `move2` objects without coercion to other movement classes. A useful overview of the wider package landscape is given by Joo *et al*. (2020).

## Data and code availability

The `move2utils` package is open-source under GPL (≥3). It is developed at https://gitlab.mpcdf.mpg.de/anenvi/r-packages/move2utils, where continuous integration runs the full `R CMD check` suite and builds the documentation site, and is mirrored for public installation at https://github.com/move2universe/move2utils. It can be installed directly from either remote:

~~~
remotes::install_github(“move2universe/move2utils”)
remotes::install_gitlab(“anenvi/r-packages/move2utils”,
                             host = “gitlab.mpcdf.mpg.de”)
~~~

A CRAN release is planned. The fisher tracking data used in the worked example are bundled with `move2` and accessible via `move2::mt_example()` (LaPoint *et al*., 2013). The code reproducing the worked example and Figure 1 is available as a package vignette.

## Acknowledgements

We thank early adopters of `move2utils` for valuable feedback on the package’s API and documentation. The Max Planck Computing and Data Facility provides infrastructure for the package’s continuous integration, documentation hosting, and benchmark sweeps.

## Use of generative AI

We used Claude Code (Anthropic; Claude Opus 4.x models, 2026) during both the development of the `move2utils` package and the preparation of this manuscript. Within the package, the tool assisted with drafting and refactoring R code, generating scaffolding for unit tests, and surfacing legacy inconsistencies across exported functions; AI-assisted code was reviewed, tested, and integrated by the authors, and is traceable through the package’s git history. During manuscript preparation, the tool assisted with drafting, editing, and grammatical correctness, and with cross-checking claims in the text against the package source code. Literature search and verification of all cited references were performed by the authors without AI assistance. The authors reviewed all AI-generated material, take full responsibility for the content of both the package and this manuscript, and confirm that no AI tool is listed as an author.

